# TYK2 inhibition enhances Treg differentiation and function while preventing Th1 and Th17 differentiation

**DOI:** 10.1101/2024.10.01.616157

**Authors:** Karoliina Tuomela, Rosa V. Garcia, Dominic A. Boardman, Pedram Tavakoli, Maria Ancheta-Schmit, Ho Pan Sham, Lihong Cheng, Mary Struthers, Brian Bressler, Bruce A. Vallance, Qihong Zhao, Megan K. Levings

## Abstract

Janus kinase (JAK) inhibitors are widely use to inhibit inflammatory cytokine signalling in autoimmune and inflammatory diseases but their effect on regulatory T cells (Tregs) is poorly characterized. We investigated the effect of JAK inhibition on human Treg differentiation, phenotype, and function using a JAK inhibitor, upadacitinib, in comparison to BMS-986202, a selective Tyrosine kinase 2 (TYK2) inhibitor. Both upadacitinib and BMS-986202 blocked the differentiation of naïve CD4^+^ T cells into Th1 and Th17 cells, but only BMS-986202 spared IL-2 signalling and Treg differentiation. BMS-986202 also increased Treg suppressive function and stability under Th1- and Th17-polarizing conditions, whereas upadacitinib significantly impaired the phenotype and viability of *ex vivo* Tregs. Analysis of lamina propria mononuclear cells from patients with inflammatory bowel disease revealed that, under Th17 polarizing conditions, BMS-986202 redirected CD4^+^ T cells towards a Treg phenotype. The Treg-sparing and enhancing properties of TYK2 inhibition suggest that TYK2 inhibitors are a promising pharmacological approach for tolerance induction.

**eTOC SUMMARY:** Tuomela et al. report that TYK2 inhibition does not affect human Treg induction from naïve CD4^+^ T cells, promotes Treg differentiation in lamina propria-derived T cells, and increases blood-derived Treg stability/function. In contrast, JAK inhibition strongly impairs Treg function.

## INTRODUCTION

Inflammatory CD4^+^ T cell subsets, including interferon (IFN)-γ-producing T helper (Th) 1 cells and interleukin (IL)-17-producing Th17 cells, are central to the pathophysiology of many autoimmune and chronic inflammatory diseases (Sun *et al*., 2023). At homeostasis and during the resolution of inflammation, the activities of inflammatory Th subsets are balanced by regulatory T cells (Tregs) (Dikiy and Rudensky, 2023). However, during inflammation and autoimmunity, the proportion and function of Tregs is often reduced, leading to a skewing of the Th/Treg balance (Dominguez-Villar and Hafler, 2018). Therefore, a major goal of disease-modifying therapies is to reduce the differentiation and function of inflammatory Th cells, thereby shifting the Th cell/Treg balance in the favour of Tregs.

Th subset differentiation is driven by the activation of cytokine signalling cascades. Upon engagement of their cognate targets, cytokine receptors recruit one or more of the four Janus kinase (JAK) family proteins (JAK1-3 and TYK2) to trigger signal transducer and activator of transcription (STAT) family proteins, ultimately altering gene expression (Hu *et al*., 2021). Disrupting this process with small molecule JAK inhibitors has emerged as potent strategy to reduce the differentiation, activation, and function of disease-associated Th subsets in autoimmune and inflammatory diseases (Hu *et al*., 2021; Shawky *et al*., 2022). Consequently, multiple JAK inhibitors have now received FDA approval and are in clinical use to manage a variety of diseases (Shawky *et al*., 2022). However, broad inhibition of JAK-STAT signalling can have significant side-effects, including serious heart-related events, cancer, blood clots and death, as highlighted by the FDA in 2021 (FDA, 2021).

An additional limitation of some JAK inhibitors is blockade of IL-2-stimulated JAK1/3 signalling. As Tregs require IL-2-induced STAT5 phosphorylation to maintain expression of FOXP3, their lineage-defining transcription factor (Burchill *et al*., 2007; Goldstein *et al*., 2016), JAK inhibition may be deleterious to tolerance induction. Accordingly, clinical trials testing JAK inhibitors in patients with rheumatoid arthritis, myeloproliferative neoplasms, or myelofibrosis reported that JAK inhibition does not improve, and often decreases, the proportion of circulating Tregs (Massa *et al*., 2014; Keohane *et al*., 2015; Parampalli Yajnanarayana *et al*., 2015; Schroeder *et al*., 2020; Meyer *et al*., 2021; Lui *et al*., 2023). Moreover, JAK inhibitors do not restore the Th17/Treg balance in rheumatoid arthritis patients (Meyer *et al*., 2021). Thus, selective therapeutic strategies that inhibit inflammatory Th populations but allow Tregs to function are lacking.

Recently, TYK2-specific inhibitors, such as deucravacitinib, have been developed to allosterically inhibit kinase activity by binding to the pseudokinase domain (Liu *et al*., 2021; Jensen *et al*., 2023). TYK2 is involved in signal transduction downstream of many cytokines, including type I interferons, IL-10, IL-12, IL-13 and IL-23 (Hu *et al*., 2021). Supporting the rationale for TYK2 inhibition in autoimmunity, several single nucleotide polymorphisms in the *TYK2* gene, which result in reduced signalling capability, are associated with protection from systemic lupus erythematosus (SLE), type 1 diabetes (T1D), multiple sclerosis (MS), rheumatoid arthritis (RA), psoriasis, and inflammatory bowel disease (IBD) (Dendrou *et al*., 2016; Pellenz *et al*., 2021; Yuan *et al*., 2023). Moreover, mice bearing a protective *TYK2* variant, *TYK2^A^*^1104^, exhibit impaired Th1 and Th17 polarization, a reduction in IL-17^+^IFN-γ^+^ T cells, and protection against experimental autoimmune encephalitis (Gorman *et al*., 2019). Clinical trials of the TYK2-specific inhibitor, deucravacitinib, showed effective disease modification in psoriasis, psoriatic arthritis, and SLE (Morand *et al*., 2024).

Although the role of TYK2 and the effect of TYK2 inhibition in inflammatory Th1 and Th17 cells is well studied, the contribution of this pathway to Treg phenotype and function has not been directly examined. Importantly, unlike JAK1 and JAK3, TYK2 is not involved in IL-2 signalling and, thus, its therapeutic inhibition has the potential to spare Treg function while inhibiting other inflammatory cytokine pathways. In this work, we investigated the effect of BMS-986202, a selective TYK2 antagonist, on primary human Tregs and compared this to upadacitinib, a JAK inhibitor that is FDA approved for use in rheumatoid arthritis, psoriatic arthritis, and ulcerative colitis (Shawky *et al*., 2022). We found that JAK inhibition completely prevented Treg differentiation and impaired the suppressive function of ex vivo Tregs. Conversely, TYK2 inhibition promoted Treg phenotype and function and protected them from trans-differentiation into inflammatory cell types. Thus, selective TYK2 inhibition represents a new approach to simultaneously control inflammatory signalling and enhance Treg-mediated regulation.

## RESULTS

### TYK2 inhibition does not affect IL-2-induced STAT5 phosphorylation

To validate BMS-986202 as a selective TYK2 antagonist (Liu *et al*., 2021), phosphoflow experiments were performed. Primary Tregs were isolated from the peripheral blood of healthy donors and treated with cytokines that rely on TYK2 activity to trigger downstream STAT phosphorylation. Assays were performed in the presence of BMS-986202 or upadacitinib, a JAK inhibitor with highest activity against JAK1 and additional low activity against JAK2/3 (Shawky *et al*., 2022).

IFN-α-induced phosphorylation of STAT1, which is mediated through JAK1/TYK2 (Fig 1A) (Hu *et al*., 2021), was significantly reduced by both BMS-986202 and upadacitinib in a dose-dependent manner (Fig 1B-C). Similarly, both compounds reduced IL-12 and IL-23-mediated STAT phosphorylation, which relies on JAK2/TYK2 activity (Fig 1D-F) (Hu *et al*., 2021).

**Figure 1.**
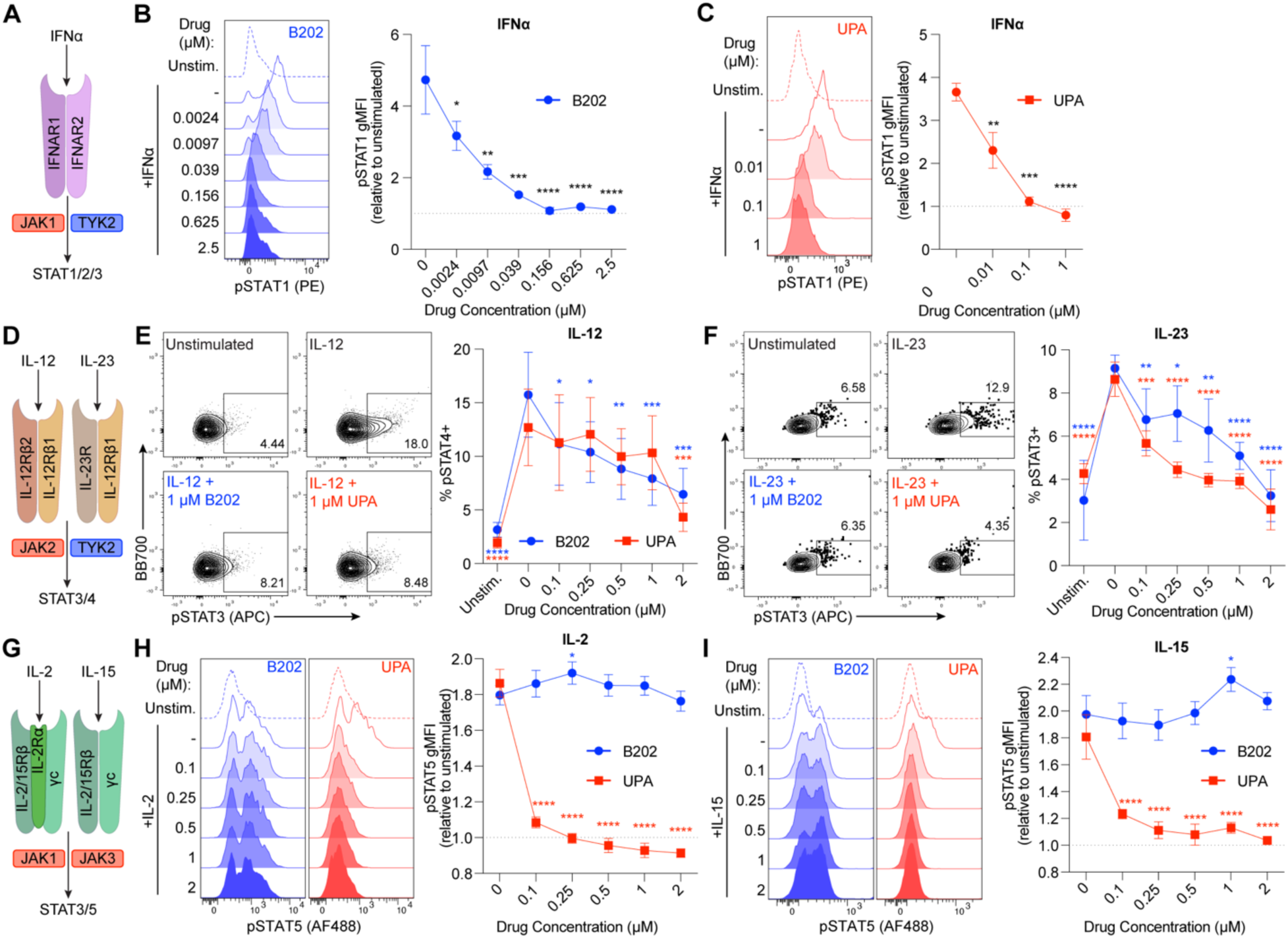
BMS-986202 inhibits TYK2-mediated cytokine signalling in Tregs. Tregs were treated with varying concentrations of BMS-986202 (B202) or upadacitinib (UPA) for 1 hour, then stimulated with cytokine for 15 min. STAT phosphorylation (pSTAT) was quantified by flow cytometry. **(A)** Schematic of IFNα signalling. **(B-C)** Representative histograms and quantification of pSTAT1 gMFI relative to unstimulated Tregs after treatment with B202 (B; n=3) or UPA (C; n=3) and stimulation with IFNα. **(D)** Schematic of IL-12/23 signalling. **(E-F)** Representative contour plots and quantification of % pSTAT3/4+ Tregs after treatment with B202 or UPA and stimulation with IL-12 (E; n=3) or IL-23 (F; n=3). **(G)** Schematic of IL-2/15 signalling. **(H-I)** Representative histograms and quantification of pSTAT5 gMFI relative to unstimulated Tregs after treatment with B202 or UPA and stimulation with IL-2 (H; n=3) or IL-15 (I; n=3). Statistical significance between DMSO (0 μM) and B202/UPA-treated cells was determined by repeated-measures one-way ANOVA with Dunnett’s multiple comparisons test (B-C) or repeated-measures two-way ANOVA with Sidak’s multiple comparisons test (E-F, H-I). * p<0.05, ** p<0.01, *** p<0.001, **** p<0.0001

To confirm the effects of BMS-986202 were TYK2-specific, we next tested whether these compounds influenced TYK2-independent cytokine signalling pathways (Fig 1G) (Hu *et al*., 2021). Upadacitinib potently prevented IL-2- and IL-15-induced STAT5 phosphorylation in Tregs (Fig 1H-I). In contrast, BMS-986202 had no impact on IL-2/15-induced pSTAT5 even at a high concentration (2 μM). Overall, TYK2 inhibition prevents IFN-α, IL-12, and IL-23 signalling but preserves IL-2/15-induced STAT5 phosphorylation in Tregs.

### TYK2 inhibition blocks the differentiation of Th1 and Th17 cells but not of Tregs

Differentiation of naïve CD4+ T cells into Th1 and Th17 subsets is induced by cytokine signalling in conjunction with TCR stimulation (Sun *et al*., 2023). Several of the cytokines involved in this process, including IL-12, IL-6, and IL-23, signal in a TYK2-dependent manner (Hu *et al*., 2021; Sun *et al*., 2023). Therefore, we assessed the ability of the TYK2 inhibitor, BMS-986202, compared to upadacitinib, to influence Th1 and Th17 differentiation of naïve human CD4+ T cells.

To induce Th17 differentiation, naïve CD4+ T cells from healthy human donors were stimulated with anti-CD3/CD28 mAbs and a cocktail of IL-23, IL-1β, TGF-β, IL-6, and anti-IL-4/IFN-γ antibodies in the presence or absence of BMS-986202 or upadacitinib (Fig 2A). After 7 days, Th17 phenotype was validated by flow cytometry, as indicated by expression of the chemokine receptors, CCR4 and CCR6, and the canonical Th17 transcription factor, RORC2 (Fig 2B). Both upadacitinib and BMS-986202 significantly reduced the expression of these Th17 markers in a dose-dependent manner when compared to control DMSO-treated cells. Intracellular IL-17A/F expression (Fig 2C) and cytokine secretion (Fig 2D) were also significantly reduced in T cells treated with either upadacitinib or BMS-986202 in a dose-dependent manner (Fig 2C). Overall, these results confirmed that Th17 differentiation is inhibited by both JAK and TYK2 inhibition.

**Figure 2.**
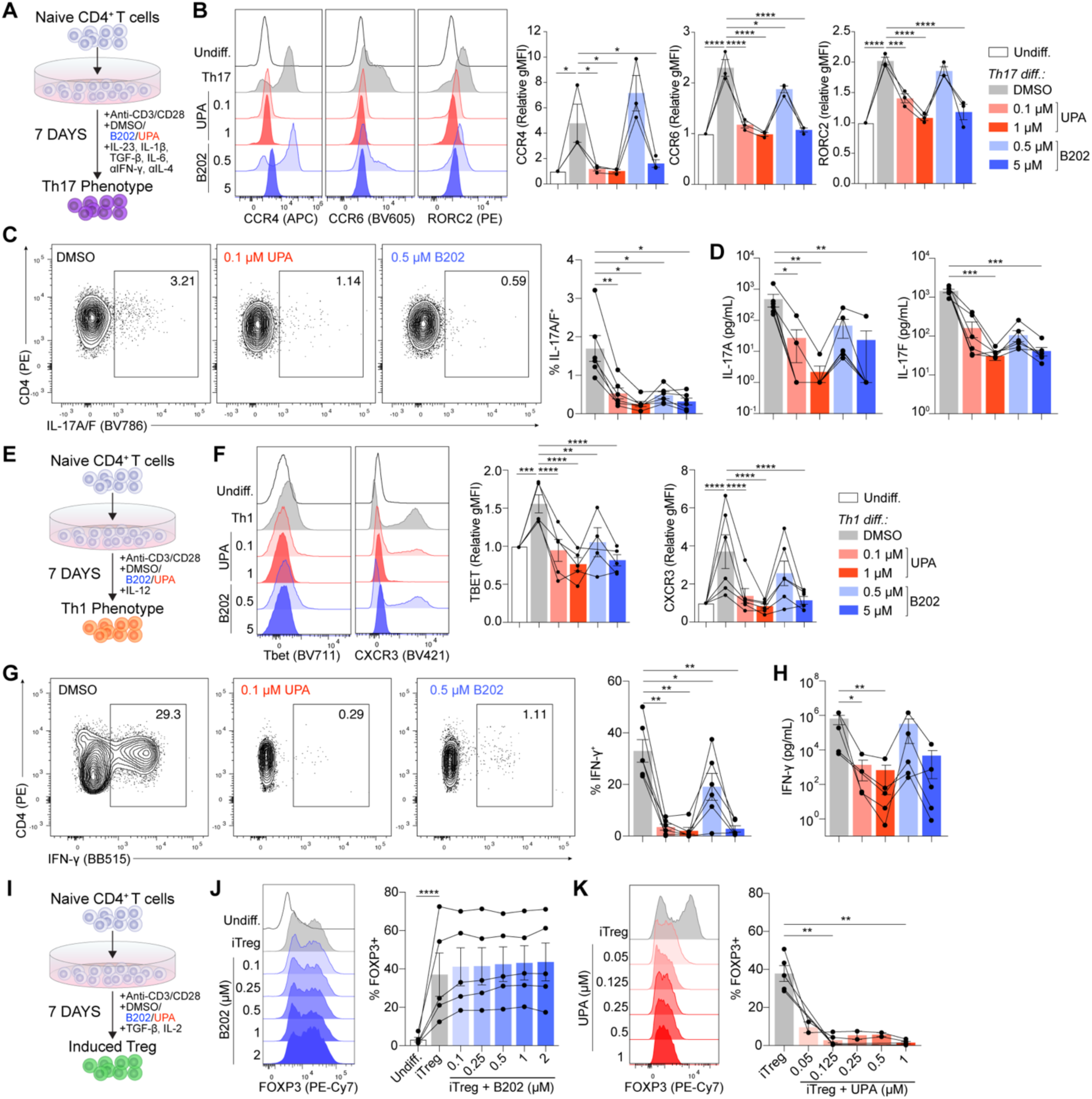
TYK2 inhibition reduces Th1 and Th17 differentiation of naïve CD4^+^ T cells but preserves Treg induction. **(A-D)** Naïve CD4^+^ T cells were stimulated with anti-CD3/CD28 for 7 days in the presence of a Th17-inducing cytokine cocktail ± BMS-986202 (B202) or upadacitinib (UPA) to induce Th17 differentiation. (A) Schematic of Th17 differentiation. (B) Representative histograms and quantification of CCR4, CCR6, and RORγt expression relative to undifferentiated CD4^+^ T cells at day 7 (n=3). (C) Representative plot and quantification of intracellular IL-17A/F expression after a 4 h stimulation with PMA/ionomycin (n=6). (D) IL-17A and IL-17F concentration in supernatant after a 24 h re-stimulation with anti-CD3/CD28 (n=6). **(E-H)** Naïve CD4^+^ T cells were stimulated with anti-CD3/CD28 for 7 days in the presence of IL-12 ± B202/UPA to induce Th1 differentiation. (E) Schematic of Th1 differentiation. (F) Representative histograms and quantification of Tbet and CXCR3 expression relative to undifferentiated CD4^+^ T cells at day 7 (n=6). (G) Representative plot and quantification of intracellular IFNγ expression after a 4 h stimulation with PMA/ionomycin (n=6). (D) IFNγ concentration in supernatant after a 24 h re-stimulation with anti-CD3/CD28 (n=6). **(I-K)** Naïve CD4^+^ T cells were stimulated with anti-CD3/CD28 for 7 days in the presence of TGFβ ± B202/ UPA to induce Treg differentiation. (I) Schematic of Treg differentiation. (J-K) Representative histograms and quantification of FOXP3 expression at day 7 with treatment with B202 (J; n=5) or UPA (K; n=2-5). Statistically significant differences compared to DMSO-treated cells were determined by a repeated-measures one-way ANOVA with Dunnett’s multiple comparisons test (B-D, F-H, J). * p<0.05, ** p<0.01, *** p<0.001, **** p<0.0001

To induce Th1 differentiation, naïve CD4+ T cells were stimulated with anti-CD3/CD28 mAbs and IL-12 in the presence of either BMS-986202 or upadactinib (Fig 2E). After 7 days, Th1-differentiated T cells upregulated TBET and CXCR3 (Fig 2F) and produced IFN-γ (Fig 2G-H). Treatment with either upadacitinib or BMS-986202 reduced the expression of TBET and CXCR3 under Th1-dfferentiating conditions in a dose-dependent manner. Similarly, intracellular IFN-γ expression and secretion were reduced by upadacitinib and BMS-986202 compared to untreated cells (Fig 2G-H).

Naïve CD4+ T cells can also be differentiated *in vitro* into Tregs via TCR stimulation in the presence of TGF-β and IL-2. Peripherally-differentiated Tregs are critical in the control of autoimmunity and inflammation (Dominguez-Villar and Hafler, 2018). As such, we compared the effects of JAK versus TYK2 inhibition on this process (Fig 2I). As anticipated, after 7 days of culture in Treg-inducing conditions, we observed a significant increase in FOXP3 expression (Fig 2J-K). However, unlike with Th1 and Th17 differentiation, TYK2 inhibition by BMS-986202 did not disrupt Treg induction at any concentration tested (Fig 2J). In contrast, upadacitinib potently prevented Treg induction at concentrations as low as 0.05 μM. Thus, although both inhibitors reduce Th1 and Th17 differentiation from naïve CD4 T cells, only TYK2 inhibition preserves Treg induction.

### TYK2 inhibition preserves the Treg phenotype and enhances their suppressive function

Having confirmed that TYK2 inhibition does not interfere with Treg differentiation from naïve CD4^+^ T cells *in vitro* (Fig 2I-K), we next sought to compare the effect of TYK2 and JAK inhibition on *ex vivo* Tregs isolated from peripheral blood. CD4^+^CD25^hi^CD127^lo^ Tregs from healthy human donors were stimulated and cultured for 7 days in the presence of IL-2 and varying concentrations of the indicated drugs (Fig 3A). Upadacitinib-mediated JAK inhibition potently reduced the viability (Fig 3B) and cell cycling (Fig 3C) of Tregs. Upadacitinib also significantly reduced the expression of the lineage-defining transcription factor, FOXP3, in a dose-dependent manner, with a lower proportion of FOXP3^+^ cells and FOXP3^+^Helios^+^ cells (Fig 3D), the latter of which are thought to represent stable thymic-derived Tregs. In contrast, BMS-986202 had no effect on the survival, proliferation, or phenotype of Tregs, even at a high concentration of 5 μM.

**Figure 3.**
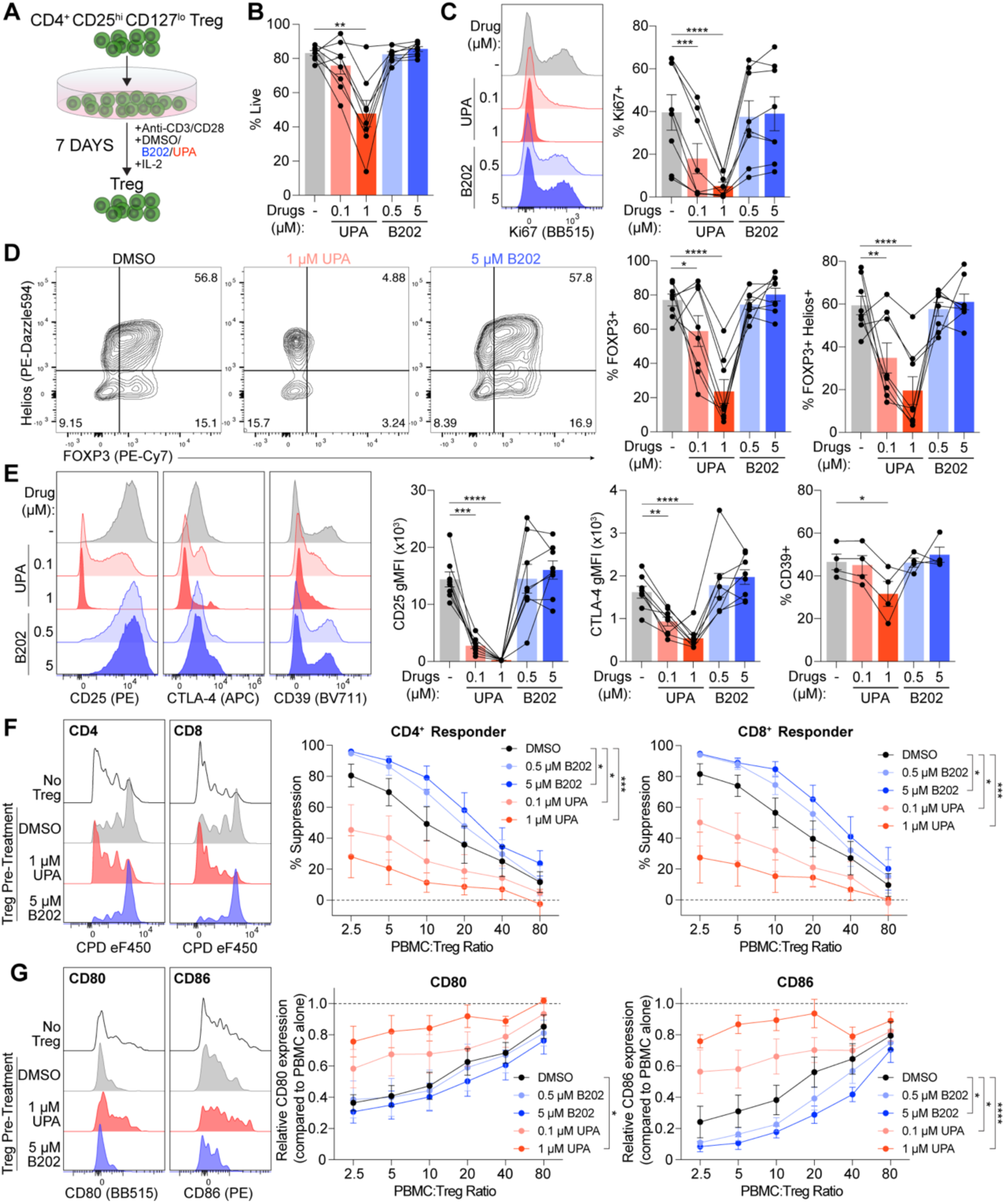
TYK2 inhibition maintains Treg phenotype and enhances suppressive function. Tregs (CD4^+^CD25^hi^CD127^lo^) were isolated from peripheral blood of healthy human donors, stimulated with anti-CD3/CD28, and cultured for 7 days in the presence of BMS-986202 (B202) or upadacitinib (UPA). **(A)** Schematic of Treg culture. **(B-E)** Treg viability (B), percentage Ki67^+^ (C), percentage FOXP3^+^/Helios^+^ (D), and CD25, CTLA-4, and CD39 expression were determined after 7 days (n=4-8). **(F-G)** Responder PBMCs were stimulated with anti-CD3/CD28 Dynabeads in the presence of varying ratios of Tregs and cultured for 96 hours. (F) Percent suppression of CD4^+^ and CD8^+^ responder T cell proliferation was determined relative to PBMCs alone (dotted line). Representative histograms and quantification shown (n=7). (G) CD80 and CD86 expression on B cells relative to PBMC alone condition (dotted line). Representative histograms and quantification shown (n=6). Statistically significant differences compared to DMSO-treated cells were determined by one-way ANOVA with Dunnett’s multiple comparisons test (B-E) or an uncorrected Fisher’s LSD test (F-G). * p<0.05, ** p<0.01, *** p<0.001, **** p<0.0001

We also observed a potent reduction in expression of the high affinity IL-2 receptor, CD25, on Tregs cultured with upadacitinib (Fig 3E). Upadacitinib also reduced expression of the Treg effector molecules, CTLA-4 and CD39 (Fig 3E). In contrast, TYK2 inhibition by BMS-986202 had no impact on CD25, CTLA-4, or CD39 expression. Expression of other markers associated with Treg function and stability, namely TIGIT and TIM-3, were also impaired by upadacitinib but not BMS-986202 (Supp Fig 1A).

We also investigated the effect of BMS-986202 on the production of cytokines by Tregs during culture. Supernatants were collected after 3 days of culture and analysed (Supp Fig 1B). Tregs produced detectable amounts of IFN-γ, IL-13, and IL-5, and very low amounts of IL-17A, IL-10, IL-22, and IL-9. Culture with BMS-986202 significantly reduced secretion of the Th1-related cytokine, IFN-γ, but increased that of the Th2-related cytokines, IL-13 and IL-5.

Given the aforementioned negative impact of upadacitinib on Treg functional marker expression, we next tested the suppressive capacity of Tregs which were cultured for 7 days in the presence of varying concentrations of upadacitinib or BMS-986202. To test suppression, PBMCs were stimulated with anti-CD3/CD28-coated beads and cultured in the absence or presence of Tregs which had been pre-treated with inhibitors for 72 hours, then washed so the assay was in the absence of any inhibitors. Consistent with the dose-dependent reduction in expression of functional molecules by upadacitinib, upadacitinib-treated Tregs were significantly less able to suppress CD4^+^ and CD8^+^ T cell proliferation (Fig 3F) and reduce expression of the co-stimulatory molecules, CD80 and CD86, on B cells (Fig 3G). Conversely, Tregs pre-treated with the TYK2 antagonist, BMS-986202, had the reverse effect, resulting in enhanced suppression of CD4^+^ and CD8^+^ T cell proliferation and CD86 expression on B cells compared to untreated Tregs (Fig 3F-G). Thus, TYK2 inhibition spares the phenotype of Tregs and promotes their suppressive function.

### TYK2 inhibition promotes Tregs stability

In inflammation and autoimmunity, Tregs can acquire a Th1- or Th17-like phenotype characterized by expression of inflammatory cytokines (Contreras-Castillo *et al*., 2024). Polarization towards these inflammatory phenotypes is driven by similar cytokine conditions as for canonical Th1 and Th17 differentiation: IL-12 for Th1 and a combination of IL-23, IL-1β, TGF-β, and IL-6 for Th17.

We first explored the effect of Th17 polarizing cytokines on the stability, phenotype, and function of peripheral blood-derived human Tregs in the presence or absence of BMS-986202 (Fig 4A). After 7 days, Th17 polarization had no impact on the expression of FOXP3 or methylation status of the Treg-specific demethylated region (TSDR), indicating that Tregs remain stable in this cytokine milieu (Fig 4B, D; Supp Fig 2A). However, Th17-polarized Tregs gained expression of canonical Th17 markers, such as RORC2, as well as increased expression and secretion of IL-17A, IL-17F, and TNF-α (Fig 4C, E-F, Supp Fig 2B-C). Culture with BMS-986202 inhibited RORC2 upregulation and reduced CCR6 expression (Fig 4C). Intracellular expression of IL-17A/F and secretion of IL-17A, IL-17F, and TNF-α were also significantly reduced by TYK2 inhibition (Fig 4E-F; Supp Fig 2C). Interestingly, BMS-986202 synergized with Th17-polarizing conditions to significantly increase CCR4 expression (Fig 4C), which marks lymph node-homing, memory-like Tregs in addition to Th17 cells (Baatar *et al*., 2007; Yuan *et al*., 2007). BMS-986202 also significantly reduced methylation of the TSDR in both control and Th17-polarizing conditions (Fig 4D).

**Figure 4.**
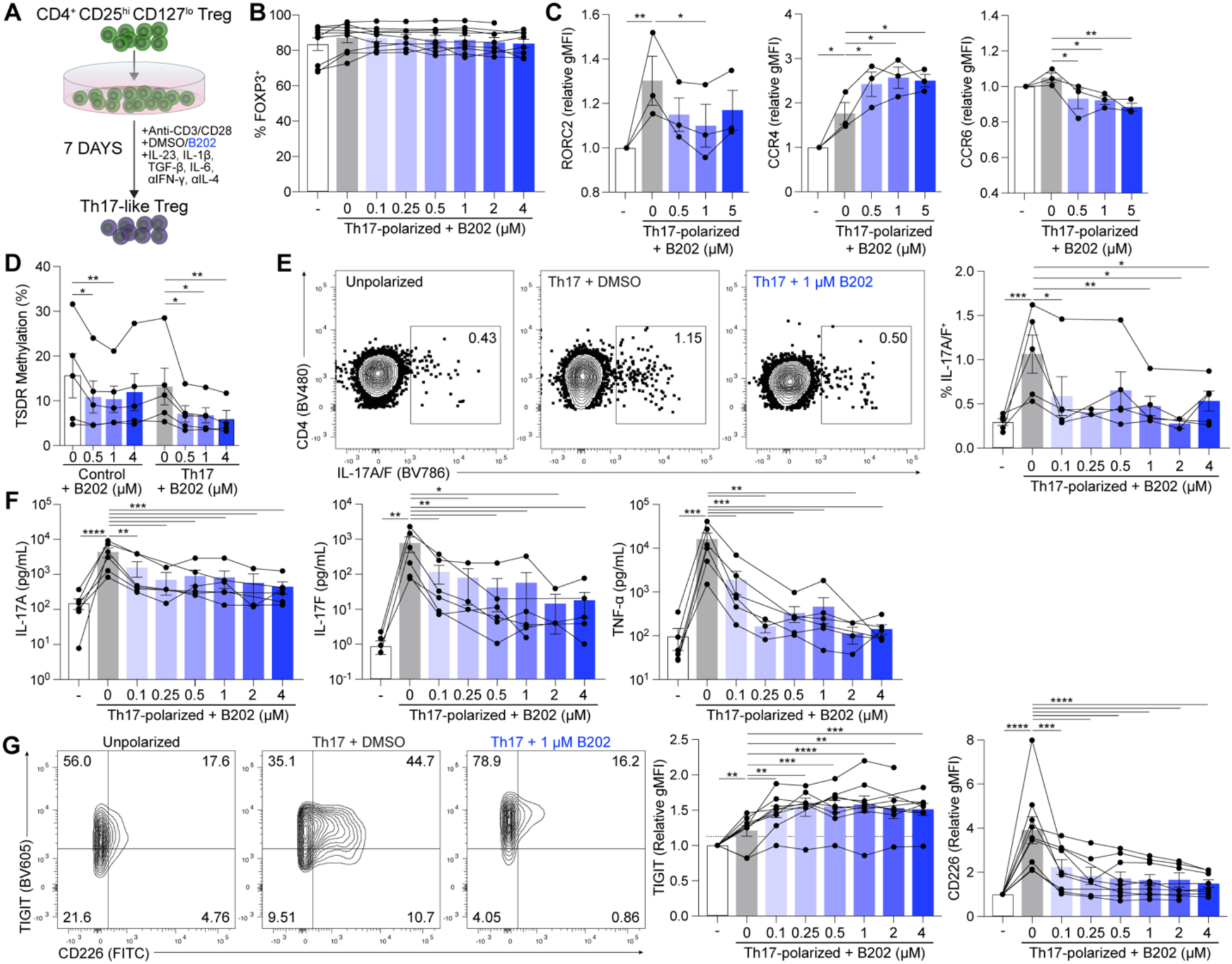
TYK2 inhibition enhances Treg stability in Th17-polarising conditions. Tregs (CD4^+^CD25^hi^CD127^lo^) were isolated from peripheral blood of healthy human donors, stimulated with anti-CD3/CD28, and cultured for 7 days in a Th17-polarizing cocktail in the presence of BMS-986202 (B202). **(A)** Schematic of polarization protocol. **(B)** The proportion of FOXP3^+^ cells was assessed after 7 days. Representative plots and quantification shown (n=9). **(C)** Expression of RORγt, CCR4, and CCR6 was determined after 7 days relative to non-polarized, control Tregs (n=3). **(D)** Methylation of the Treg Specific Demethylated Region (TSDR) after 7 days of culture (n=5). **(E)** Representative plot and quantification of intracellular IL-17A/F expression after a 4 h stimulation with PMA/ionomycin (n=5). **(F)** IL-17A, IL-17F, and TNFα concentration in supernatant after a 24 h re-stimulation with anti-CD3/CD28 (n=6). **(G)** Representative figures and quantification of TIGIT and CD226 expression on Tregs on day 7 (n=9). Statistically significant differences compared to DMSO-treated cells were determined by a repeated-measures one-way ANOVA with Dunnett’s multiple comparisons test (B-G). * p<0.05, ** p<0.01, *** p<0.001, **** p<0.0001

The balance of the co-inhibitory receptor TIGIT and the co-stimulatory receptor CD226 has been identified as a regulator of Treg stability under inflammatory conditions (Fuhrman *et al*., 2015; Fourcade *et al*., 2018; Thirawatananond *et al*., 2023). We found that Th17 polarization strongly upregulated CD226 on Tregs, and this was blocked in a dose-dependent manner by BMS-986202 (Fig 4G). Although TIGIT expression was marginally increased in Th17-polarizing conditions, BMS-986202 led to a further upregulation (Fig 4G). Thus, TYK2 inhibition blocks the polarization of human Tregs towards a more inflammatory phenotype characterised by RORC2, IL-17A, IL-17F, TNF-α, and CD226 expression.

Next, we investigated the effect TYK2 inhibition on IL-12-induced Th1 polarization of Tregs (Fig 5A). IL-12 did not affect FOXP3 expression (Fig 5B) but was associated with increased intracellular expression and secretion of IFN-γ (Fig 5C-D), indicating that Tregs are polarized towards a Th1-like phenotype. Treatment with BMS-986202 significantly reduced the proportion of IFN-γ^+^ cells and overall IFN-γ secretion in a dose-dependent manner (Fig 5C-D). Interestingly, IL-12 also increased the proportion IFN-γ^+^ IL-17A/F^+^ Tregs, a particularly proinflammatory subtype of cell, but this was significantly reduced by BMS-986202 (Fig 5E). In contrast to Th17 polarization, Th1 polarization alone did not affect the expression of CD226 or TIGIT on Tregs, but additional treatment with BMS-986202 did increase TIGIT expression (Fig 5F). Overall, treatment of Tregs with BMS-986202 promotes lineage stability by preventing Th1 and Th17 polarization and gain of inflammatory cytokine expression and enhancing the epigenetic hallmark of Tregs, namely de-methylation of the TSDR region.

**Figure 5.**
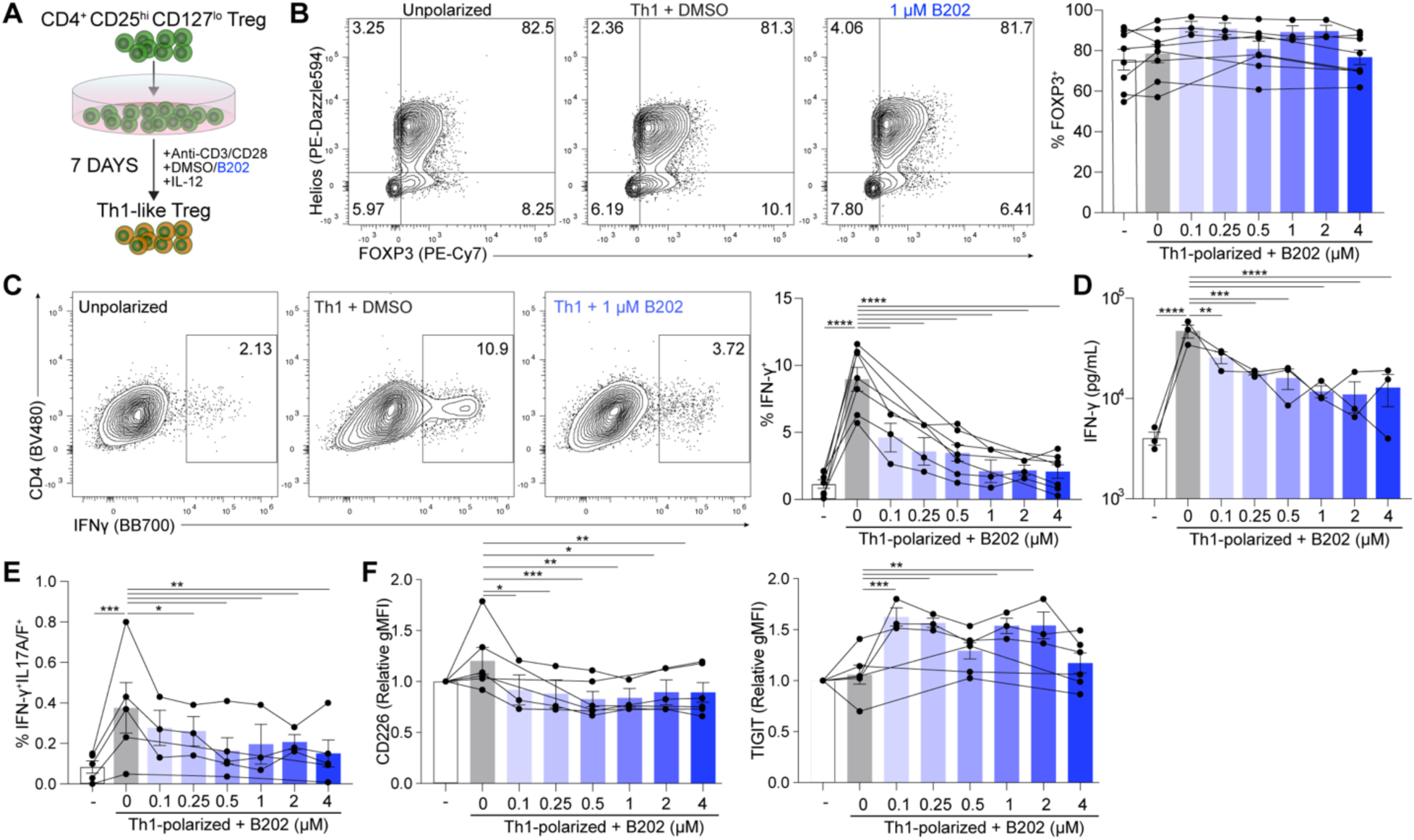
TYK2 inhibition enhances Treg stability in Th1-polarizing conditions. Tregs (CD4^+^CD25^hi^CD127^lo^) were isolated from peripheral blood of healthy human donors, stimulated with anti-CD3/CD28, and cultured for 7 days with IL-12 and in the presence of BMS-986202 (B202). **(A)** Schematic of polarization protocol. **(B)** The proportion of FOXP3^+^ cells was assessed after 7 days. Representative plots and quantification shown (n=3-8). **(C)** Representative plots and quantification of intracellular IFNγ expression after a 4 h stimulation with PMA/ionomycin (n=3-7). **(D)** IFNγ concentration in supernatant after a 24 h re-stimulation with anti-CD3/CD28 tetramer (n=3). **(E)** Proportion of Tregs with intracellular expression of IFNγ^+^IL-17A/F^+^ Tregs after a 4 h stimulation with PMA/ionomycin (n=3-5). **(F)** Surface expression of CD226 and TIGIT on Tregs on day 7 (n=3-6). Statistical differences between DMSO and other conditions were determined by a repeated-measures one-way ANOVA with Dunnett’s multiple comparisons test (B-G). * p<0.05, ** p<0.01, *** p<0.001, **** p<0.0001

### TYK2 inhibition redirects gut-infiltrating T cells from Th17 to Treg differentiation

T cell phenotypes are affected by tissue localisation and disease state (Masopust and Soerens, 2019). Thus, we investigated the effect of TYK2 inhibition on T cells derived from the lamina propria of healthy individuals and patients with IBD, a disease that is frequently treated with JAK inhibitors (Hu *et al*., 2021; Shawky *et al*., 2022). Lamina propria mononuclear cells (LPMCs) were isolated from colonic biopsies of IBD or non-IBD patients, stimulated via CD3/CD28, and cultured for 7 days in the presence of Th17-polarising cytokines with or without BMS-986202 (Fig 6A). Supernatants were collected after 3 days of culture and the cytokine signature was investigated. LPMCs cultured in Th17 conditions were characterised by secretion of IL-17A, IL-17F, and TNF-α (Fig 6B; Supp Fig 3A). However, secretion of these cytokines was significantly diminished by BMS-986202. BMS-986202 additionally reduced IFN-γ and IL-10 secretion. Interestingly, production of IL-22, which is involved in the maintenance of gut homeostasis (Keir *et al*., 2020), as well as IL-13 were increased by TYK2 inhibition, collectively indicating an inhibition of Th17 immune responses. No difference in response between IBD and non-IBD patients was observed.

**Figure 6.**
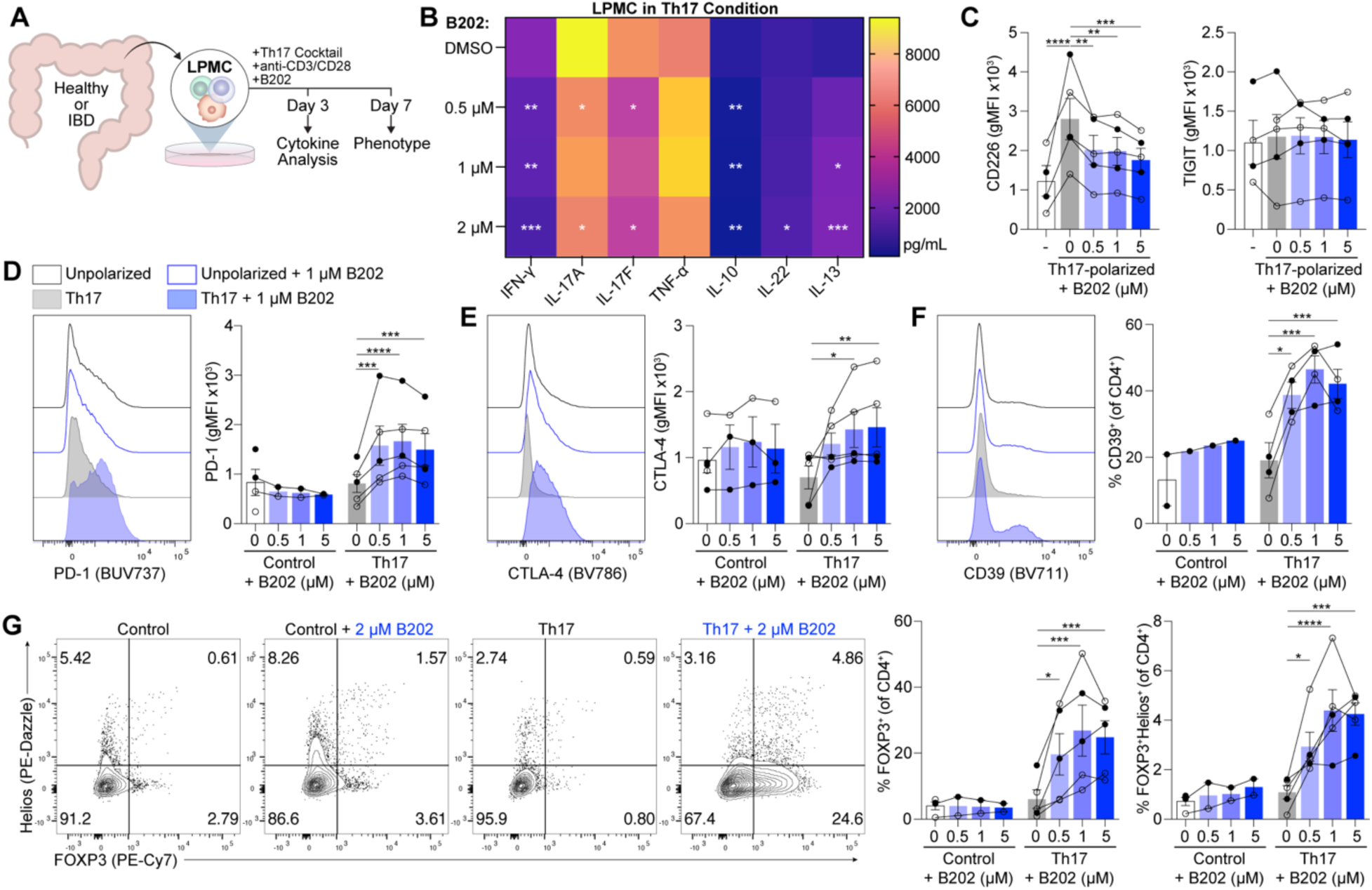
TYK2 inhibition redirects Th17-polarized LPMCs towards a regulatory phenotype. Lamina propria mononuclear cells (LPMCs) were isolated from colon biopsies of healthy or IBD patients, stimulated with anti-CD3/CD28 tetramer in the presence of a Th17 cytokine cocktail and BMS-986202 (B202), and cultured for 7 days. **(A)** Schematic of LPMC culture. **(B)** Heatmap of cytokine content in LPMC supernatant collected after 3 days of culture. **(C)** Surface expression of CD226 and TIGIT on LPMC CD4^+^ T cells. **(D-F)** Representative histograms and quantification of PD-1 (D), CTLA-4 (E), and CD39 (F) expression on LPMC CD4^+^ T cells. **(G)** Representative plots and quantification of FOXP3^+^ and FOXP3^+^Helios^+^ cells as a proportion of LPMC CD4^+^ T cells. Open circles (o) represent healthy patients. Closed circles (•) represent IBD patients. Statistically significant differences compared to DMSO-treated cells were determined by repeated-measures one-way ANOVA with Dunnett’s multiple comparisons test. * p<0.05, ** p<0.01, *** p<0.001, **** p<0.0001

We also examined the phenotype of CD4^+^ T cells within LPMCs after 7 days of culture by flow cytometry. Consistent with previous observations, Th17-polarising conditions drove increased expression of CD226 on LPMC CD4^+^ T cells from both IBD and non-IBD donors, but this was significantly reduced by BMS-986202, indicating a reduction in inflammatory activation (Fig 6C, Supp Fig 3B). However, TIGIT expression was not affected in any condition (Fig 6C; Supp Fig 3B).

Further supporting an overall shift towards a less inflammatory and more immunosuppressed phenotype, BMS-986202 increased the expression of immune checkpoint receptors, PD-1 and CTLA-4, as well as the immunoregulatory ectoenzyme, CD39, on CD4+ T cells (Fig 6D-F). Interestingly, this effect was only observed under Th17 conditions and not in control conditions without exogenous cytokines, suggesting that BMS-986202 acts to redirect T cells from an inflammatory phenotype towards a suppressed state. We did not observe significant upregulation of RORC2 in the Th17 condition, but CCR4 and CCR6 were both upregulated (Supp Fig 3C). As observed in blood-derived Tregs, BMS-986202 synergized with Th17 conditions to increase CCR4 expression, whereas CCR6 and RORC2 were not affected.

We next assessed whether the proportion of Tregs within the LPMC population was influenced by TYK2 inhibition. Surprisingly, we found that in Th17 conditions, BMS-986202 promoted a significant, dose-dependent increase in the proportion of FOXP3^+^ and FOXP3^+^Helios^+^ cells (Fig 6G). This was not observed in control (i.e. non-Th17) conditions with the addition of BMS-986202, indicating that, in LMPCs, TYK2 inhibition redirects CD4^+^ T cells from a Th17 lineage towards a Treg phenotype. However, TYK2 inhibition alone in the absence of cytokines does not drive Treg differentiation.

In parallel, we investigated the effect of TYK2 inhibition on LPMCs cultured in control and Th1 environments. Consistent with the effect of BMS-986202 on Th17-polarized LPMCs (Fig 6B), IFN-γ production was significantly reduced by TYK2 inhibition in both control and Th1 conditions (Supp Fig 4A-B). However, we did not observe any effect of Th1 polarization or BMS-986202 treatment on TIGIT/CD226 expression, Treg proportion, or expression of CTLA-4 and PD-1 (Supp Fig C-E). Overall, TYK2 inhibition shifts the phenotype of gut-infiltrating T cells in a Th17 environment from inflammation towards regulation, as evidenced by increased Treg numbers and immune checkpoint receptor expression

## DISCUSSION

Finding therapies that block inflammation whilst sparing, or even enhancing, Treg function is a long sought after goal for numerous autoimmune and autoinflammatory disorders. TYK2 inhibition using BMS-986202 surprisingly enhanced Treg phenotype, epigenetic stability, and function, while simultaneously preventing the differentiation of naïve CD4^+^ T cells and Tregs into Th1- and Th17-like cells. In LPMCs from IBD and non-IBD donors cultured in Th17 cytokines, TYK2 inhibition promoted an overall shift from an inflammatory to a regulatory state, as evidenced by reduced cytokine production, upregulation of immune checkpoint receptors on CD4^+^ T cells, and an increased proportion of Tregs. In contrast, whilst JAK inhibition suppressed inflammatory Th differentiation, it also potently repressed the survival, function and differentiation of Tregs. These data suggest that TYK2 inhibition could be a promising strategy to promote tolerance induction and support long-lasting re-programming of autoimmune/autoinflammatory immune responses.

Autoimmune and inflammatory diseases, such as IBD, are marked by an increase in T cell activation and reduction in Treg numbers and suppressive function (Dominguez-Villar and Hafler, 2018; Guan, 2019). We observed that TYK2 inhibition in LPMCs from both IBD and non-IBD cultured in control, Th1, or Th17 conditions resulted in a reduction in inflammatory cytokine production. However, TYK2 inhibition in LPMCs cultured in Th17 conditions uniquely shifted CD4^+^ T cells from an inflammatory to a regulated phenotype, as evidenced by reduced expression of CD226 and increased expression of PD-1, CTLA-4, and CD39 on CD4^+^ T cells. Moreover, TYK2 inhibition redirected T cells from a Th17 towards a Treg phenotype, marked by a reduction in IL-17 secretion and an increase in FOXP3^+^ cells. This induction of Tregs likely resulted from the inhibition of IL-23, a TYK2-dependent cytokine involved in Th17 differentiation and inhibition of Treg induction (Izcue *et al*., 2008; Hu *et al*., 2021). Thus, upon TYK2 blockade in the Th17 cytokine context, TGF-β drives Treg instead of Th17 differentiation (Moreau *et al*., 2022), leading to a shift in the Th17/Treg balance towards Tregs. This shift of CD4^+^ T cells towards a regulated phenotype was not observed in LPMCs cultured without exogenous cytokines or with IL-12 to induce Th1 polarization, possibly because exogenous TGF-β was not present, suggesting that TYK2 inhibition may be particularly potent in diseases with Th17 involvement.

Under inflammatory conditions, Tregs can exhibit plasticity to gain Th1-like or Th17-like properties, characterised by secretion of IFN-γ and IL-17, respectively (Contreras-Castillo *et al*., 2024). Although the function of Th-polarized Tregs is not fully understood, they are upregulated in disease and often exhibit reduced suppressive function in addition to increased inflammatory cytokine secretion (Hovhannisyan *et al*., 2011; McClymont *et al*., 2011; Duhen *et al*., 2012; Komatsu *et al*., 2014; Contreras-Castillo *et al*., 2024). We found that Th1- and Th17-polarized Tregs secreted IFN-γ and IL-17, respectively, and this was prevented by TYK2 inhibition. Moreover, TYK2 inhibition in Th17-polarized Tregs blocked the upregulation of CD226, which has been shown to mediate Treg instability and dysfunction by interfering with TIGIT signalling (Fuhrman *et al*., 2015; Fourcade *et al*., 2018; Sato *et al*., 2021; Thirawatananond *et al*., 2023). Thus, TYK2 inhibition enhances Treg stability in inflammatory conditions as exhibited by reduced expression of inflammatory cytokines and a shift in the CD226/TIGIT balance.

Surprisingly, we found that TYK2 inhibition improved Treg suppressive function even in the absence of exogenous inflammatory cytokines. Tregs can suppress effector T cell proliferation and activation by a number of mechanisms, including competition for IL-2 by CD25 expression, CTLA-4-mediated downregulation of CD80/86 on APCs, anti-inflammatory cytokine production, and adenosine production by CD39 ectoenzymes (Dikiy and Rudensky, 2023). Unlike JAK inhibition, TYK2 inhibition preserved expression of CD25, CTLA-4, and CD39, and had no deleterious effect on Treg viability or proliferation. TYK2 inhibition also significantly increased demethylation of the TSDR region, suggesting that the enhanced suppressive function maybe related to improved Treg lineage stability. However, we unexpectedly observed that TYK2 inhibition increased IL-5 and IL-13 production and CCR4 expression by Tregs, which implies polarisation towards Th2 phenotype. Little is known about the role of Th2-like Tregs, but they retain suppressive function and appear to be enriched at tumour sites through CCL17/22-mediated chemotaxis (Halim *et al*., 2017). Further work is needed to define the role of this population and understand the drivers of Th2 polarization in Tregs.

In contrast to TYK2 inhibition, JAK inhibition had a profoundly deleterious effect on the differentiation of Tregs from naïve T cells, as well as the phenotype and function of blood-derived Tregs. This agrees with recent findings by Satoh-Kanda et al. that BMS-986202 does not impact Treg differentiation, whereas other Jak inhibitors abolish induction (Satoh-Kanda *et al*., 2024). Similarly, TYK2-KO mouse T cells exhibit normal TGF-β-induced Treg differentiation (Ishizaki *et al*., 2011). Together, these data demonstrate that TYK2 inhibition is likely to be compatible with Treg-dependent mechanisms of tolerance induction in autoimmunity, whereas JAK inhibition is not. Furthermore, this presents an opportunity for combination of TYK2 inhibitors with other Treg-promoting therapies, such as adoptive Treg therapy (Tuomela, Salim and Levings, 2023).

In addition to BMS-986202, other TYK2 inhibitors have been developed, including deucravacitinib (BMS-986165), brepocitinib (PF-06700841), and PF-06826647. Neither brepocitinib nor PF-06826647 are fully TYK2-specific, with additional inhibitory effects at clinically relevant concentrations on JAK1 and JAK2, respectively (Fensome *et al*., 2018; Gerstenberger *et al*., 2020). The effect of non-specific TYK2 inhibitors on Tregs has not been investigated in the context of clinical trials, but a recent study demonstrated that brepocitinib is not permissive to Treg differentiation (Satoh-Kanda *et al*., 2024). Although it has not been directly investigated, PF-06826647 may spare Tregs as it primary targets TYK2 and JAK2, which are not involved in IL-2 signalling (Gerstenberger *et al*., 2020). In contrast to these inhibitors, BMS-986202 and deucravacitinib are allosteric inhibitors targeting the pseudokinase domain of TYK2, leading to far better selectivity (Wrobleski *et al*., 2019; Liu *et al*., 2021). Deucravacitinib was initially tested and approved for use in psoriasis (Armstrong *et al*., 2023; Strober *et al*., 2023), but clinical trials have since expanded its use to SLE and psoriatic arthritis with positive results (Mease *et al*., 2022; Morand *et al*., 2023). However, deucravacitinib failed to meet the primary efficacy endpoint in an ulcerative colitis trial (Danese *et al*., 2022), despite initial positive findings in a mouse colitis model (Burke *et al*., 2019). This lack of efficacy may result from the pleiotropic role of Th17 cells in IBD pathogenesis, with reports that IL-17 inhibition can precipitate or exacerbate IBD (Deng *et al*., 2023). Indeed, our data demonstrate that TYK2 inhibition reduces Th17 polarisation and activity in LPMCs. It is possible that TYK2 inhibition could be more effective in diseases in which Th17 cells have a clear pathogenic role, such as psoriasis and MS (Schnell, Littman and Kuchroo, 2023). Other diseases, such as type 1 diabetes, in which TYK2 has been shown to contribute to autoimmune attack are also promising targets (Chandra *et al*., 2022; Mine *et al*., 2024).

In summary, we observed that TYK2 inhibition preserves Treg phenotype, enhances suppressive function, and shifts the Th17/Treg balance towards a regulated phenotype. In contrast, the JAK inhibitor, upadacitinib, had a profoundly negative impact on Tregs and Treg differentiation. Overall, TYK2 inhibition has a targeted effect on inflammatory pathways, whilst preserving the regulatory component of the immune system. Therefore, improving Treg function via TYK2 inhibitors will likely contribute to long-term re-establishment of immune homeostasis and, as a result, promote durable remission in chronic inflammatory and autoimmune diseases.

## MATERIALS AND METHODS

### PBMC and T cell isolation

Leukopaks from healthy donors were obtained from StemCell Technologies (Vancouver, Canada). All samples were obtained with informed consent and ethical approval (The University of British Columbia Clinical Research Ethics Board number H15-02826). PBMCs were isolated by density gradient centrifugation with Lymphoprep Density Gradient Medium (STEMCELL Technologies). To isolate CD4+ T cells, the RosetteSep human CD4+ T cell enrichment cocktail (STEMCELL Technologies) was used prior to density gradient centrifugation. To isolate Tregs from CD4+ T cells, CD25 enrichment was carried out (CD25 Microbeads II; Miltenyi Biotec). Total Tregs (CD4+CD127loCD25hi) were sorted from the CD25+ fraction using a MoFlo Astrios (Beckman Coulter) or FACSAria Fusion (BD Biosciences). To isolate naïve CD4+ T cells from total CD4+ T cells, CD45RA enrichment was carried out using the Easy Sep Naïve CD4 Isolation Kit (STEMCELL Technologies) prior to sorting for CD4+CD25-CD127+CD45RA+CD45RO-CD62L+ cells. T cells were maintained in Immunocult-XF media (STEMCELL Technologies) supplemented with 1% penicillin-streptomycin (Gibco) unless otherwise described.

### Th cell differentiation/polarization

Tregs or naïve CD4+ T cells were stimulated with ImmunoCult Human CD3/CD28 T cell Activator (STEMCELL Technologies) in the presence of Th1- (10 ng/mL IL-12 [BD Biosciences]), Th17- (10 ng/mL IL-1β [STEMCELL Technologies], 10 ng/mL IL-6 [STEMCELL Technologies], 10 ng/mL TGFβ [R&D Systems], 20 ng/mL IL-23 [PeproTech], 5 μg/mL anti-IFN-γ, 5 μg/mL anti-IL-4), or Treg-inducing cytokines (5 ng/mL TGFβ), as previously described (Boardman *et al*., 2020). For all conventional CD4^+^ T cell culture, 100 U/mL IL-2 (Proleukin) was included. For all Treg culture, 1000 U/mL IL-2 was included. Media was replenished at day 3 and day 5 and cells were analysed on day 7.

### Flow cytometry

Cells were washed and stained in PBS supplemented with 1% bovine serum (HyClone) with fluorescent antibodies against surface markers and fixable viability dye (FVD eFluor780; Thermo Fisher Scientific) for 30 min at 4°C. All surface antibodies are described in Supplementary Table 1. Cells were fixed and permeabilized using eBioscience FOXP3/Transcription Factor Staining Buffer Set (Invitrogen) before staining for intracellular markers. Intracellular antibodies are described in Supplementary Table 1. Data were acquired using a BD LSRFortessa or BD FACSymphony A1 or A5, and analysed using FlowJo V10.10 Software (all BD Life Sciences).

### Cytokine Stimulation and Phosphoflow Cytometry

For IL-12 and IL-23 stimulation, Tregs were pre-stimulated with CD3/CD28 T cell Activators in the presence of 1000 U/mL IL-2 for 3 days. For all other cytokines, Tregs were used immediately upon isolation. Tregs were rested overnight in Immunocult-XF media. Tregs were treated with upadacitinib or BMS-986202 for 1 hour before stimulation with IL-12 (50 ng/mL; BD Biosciences), IL-23 (50 ng/mL; PeproTech), IFN-α (40,000 U/mL; STEMCELL Technologies), IL-2 (100 U/mL), or IL-15 (25 ng/mL; STEMCELL Technologies) for 15 min. Cells were then fixed and permeabilized using the Transcription Factor Phospho Buffer Set (BD Pharmingen) according to the manufacturer’s protocol. Cells were stained using anti-pSTAT1, anti-pSTAT3, anti-pSTAT4, or anti-pSTAT5 (Supplementary Table 1) depending on the stimulating cytokine. Samples were quantified by flow cytometry (BD LSRFortessa or BD FACSymphony) and analysed using FlowJo V10.10 Software (BD Life Sciences).

### Cytokine analysis

To quantify cytokine secretion, cell-free supernatant was collected from cultures 3 days after initial stimulation or 24 hours after re-stimulation with CD3/CD28 T cell Activators. Cytokines were quantified using the LEGENDplex 12-plex Human Th Cytokine Panel (BioLegend) according to the manufacturer’s protocol and samples were analysed using a BD FACS Symphony A1 (BD Life Sciences) flow cytometer. For intracellular cytokine staining, 2×10^5^ cells were stimulated using phorbol 12-myristate 13-acetate (PMA; 10 ng/mL; Sigma-Aldrich) and ionomycin (500 ng/mL; Sigma-Aldrich) for 4 hours at 37°C in the presence of brefeldin (10 μg/mL; Sigma-Aldrich). Cells were stained for surface and intracellular proteins as described above (Supplementary Table 1).

### TSDR analysis

DNA was isolated from cell pellets and bisulphite converted using the EZ Direct Kit (Zymo Research). The TSDR was PCR-amplified using the AllTaq PCR Kit (Qiagen) using forward (AGAAATTTGTGGGGTGGGGTAT) and biotinylated reverse (ATCTACATCTAAACCCTATTATCACAACC) primers, then run on a W96 MD pyrosequencing system (Qiagen) with a sequencing primer (AGAAATTTGTGGGGTGGG). Analysis of cytosine versus thymine incorporation at 7 CpG sites in the TSDR was carried out using Pyro Q-CpG software (Biotage).

### Treg suppression assay

Suppression assays were carried out in Immunocult-XF supplemented with 1% penicillin-streptomycin and 5% human serum (NorthBio). PBMCs were labelled with CPD eFluor™450 (Thermo Fisher Scientific) and Tregs were labelled with CPD eFluor™670 (Thermo Fisher Scientific). 1×10^5^ PBMCs were cultured with serially diluted numbers of Tregs in the presence of anti-CD3/CD28 Human T-Activator Dynabeads™ (1:8 bead:PBMC; Thermo Fisher Scientific) for 3 days. Cells were stained and analysed by flow cytometry. The division index of responder CD4+ and CD8+ T cells was calculated using the FlowJo Proliferation tool. Suppression of proliferation was calculated as previously described (McMurchy and Levings, 2012) relative to PBMCs cultured alone. Expression of CD80 and CD86 on CD19+CD3-B cells was also measured by flow cytometry and quantified relative to PBMC alone.

### LPMC isolation and culture

Intestinal biopsies were obtained during routine colonoscopy procedures from patients with or without IBD. All samples were obtained with informed consent and ethical approval (The University of British Columbia Clinical Research Ethics Board number H15-02826). Clinical data were acquired from 3 non-IBD (mean age = 46.0 ± 4.4) and 9 IBD (mean age = 36.6 ± 3.7) patients (Supplementary Table 2). Biopsies were digested for 1 hour with Collagenase Type VIII (Sigma-Aldrich) and DNAse (STEMCELL Technologies) prior to filtering through a 100 μm mesh. CD45 selection was carried out using Human CD45 Microbeads (Miltenyi Biotec) according to the manufacturer’s protocol. CD45+ cells were stimulated with CD3/CD28 T cell Activators and cultured for 7 days in Immunocult-XF media in the presence of no cytokines or polarising conditions for Th1 (10 ng/mL IL-12 [BD Biosciences]) or Th17 (10 ng/mL IL-1β, 10 ng/mL IL-6, 10 ng/mL TGF-β, 20 ng/mL IL-23, 5 μg/mL anti-IFN-γ, 5 μg/mL anti-IL-4) cells.

### Statistics

Data were analysed using GraphPad Prism 10.2.3 (GraphPad Software, Boston, Massachusetts USA). Figures show mean ± SEM, with individual donors indicated by points and connecting lines. Statistical significance was determined as described in figure legends. Statistically significant differences were defined as P < 0.05 (*), P < 0.01 (**), and P < 0.001 (***), p <0.0001 (****).

## ACKNOWLEDGEMENTS

We thank Dr. Lisa Xu for her assistance with fluorescence-activated cell sorting, and Jana Gillies for performing the TSDR methylation analysis. KT was supported by a fellowship from the Canadian Institute for Health Research and Canucks for Kids Fund Childhood Diabetes Laboratories Postdoctoral Award. DAB was supported by fellowships from the Canadian Institute for Health Research and Health Research BC. BAV holds the CH.I.L.D. Foundation Chair in Pediatric Gastroenterology. MKL is a Canada Research Chair in Engineered Immune Tolerance and receives a Scientist Salary Award from the BC Children’s Hospital Research Institute.

## REFERENCES

Armstrong, A.W. et al. (2023) ‘Deucravacitinib versus placebo and apremilast in moderate to severe plaque psoriasis: Efficacy and safety results from the 52-week, randomized, double-blinded, placebo-controlled phase 3 POETYK PSO-1 trial’, Journal of the American Academy of Dermatology, 88(1), pp. 29–39. Available at: 10.1016/j.jaad.2022.07.002.

Baatar, D. et al. (2007) ‘Human Peripheral Blood T Regulatory Cells (Tregs), Functionally Primed CCR4+ Tregs and Unprimed CCR4− Tregs, Regulate Effector T Cells Using FasL1’, The Journal of Immunology, 178(8), pp. 4891–4900. Available at: 10.4049/jimmunol.178.8.4891.

Boardman, D.A. et al. (2020) ‘Pharmacological inhibition of RORC2 enhances human Th17-Treg stability and function’, European Journal of Immunology, 50(9), pp. 1400–1411. Available at: 10.1002/eji.201948435.

Burchill, M.A. et al. (2007) ‘IL-2 Receptor β-Dependent STAT5 Activation Is Required for the Development of Foxp3+ Regulatory T Cells1’, The Journal of Immunology, 178(1), pp. 280–290. Available at: 10.4049/jimmunol.178.1.280.

Burke, J.R. et al. (2019) ‘Autoimmune pathways in mice and humans are blocked by pharmacological stabilization of the TYK2 pseudokinase domain’, Science Translational Medicine, 11(502), p. eaaw1736. Available at: 10.1126/scitranslmed.aaw1736.

Chandra, V. et al. (2022) ‘The type 1 diabetes gene TYK2 regulates β-cell development and its responses to interferon-α’, Nature Communications, 13(1), p. 6363. Available at: 10.1038/s41467-022-34069-z.

Contreras-Castillo, E. et al. (2024) ‘Stability and plasticity of regulatory T cells in health and disease’, Journal of Leukocyte Biology, p. qiae049. Available at: 10.1093/jleuko/qiae049.

Danese, S. et al. (2022) ‘DOP42 Efficacy and safety of deucravacitinib, an oral, selective tyrosine kinase 2 inhibitor, in patients with moderately-to-severely active Ulcerative Colitis: 12-week results from the Phase 2 LATTICE-UC study’, Journal of Crohn’s and Colitis, 16(Supplement_1), pp. i091–i092. Available at: 10.1093/ecco-jcc/jjab232.081.

Dendrou, C.A. et al. (2016) ‘Resolving TYK2 locus genotype-to-phenotype differences in autoimmunity’, Science Translational Medicine, 8(363), pp. 363ra149–363ra149. Available at: 10.1126/scitranslmed.aag1974.

Deng, Z. et al. (2023) ‘IL-17 inhibitor-associated inflammatory bowel disease: A study based on literature and database analysis’, Frontiers in Pharmacology, 14, p. 1124628. Available at: 10.3389/fphar.2023.1124628.

Dikiy, S. and Rudensky, A.Y. (2023) ‘Principles of regulatory T cell function’, Immunity, 56(2), pp. 240–255. Available at: 10.1016/j.immuni.2023.01.004.

Dominguez-Villar, M. and Hafler, D.A. (2018) ‘Regulatory T cells in autoimmune disease’, Nature Immunology, 19(7), pp. 665–673. Available at: 10.1038/s41590-018-0120-4.

Duhen, T. et al. (2012) ‘Functionally distinct subsets of human FOXP3+ Treg cells that phenotypically mirror effector Th cells’, Blood, 119(19), pp. 4430–4440. Available at: 10.1182/blood-2011-11-392324.

FDA (2021) FDA requires warnings about increased risk of serious heart-related events, cancer, blood clots, and death for JAK inhibitors that treat certain chronic inflammatory conditions. FDA. Available at: https://www.fda.gov/drugs/drug-safety-and-availability/fda-requires-warnings-about-increased-risk-serious-heart-related-events-cancer-blood-clots-and-death (Accessed: 29 August 2024).

Fensome, A. et al. (2018) ‘Dual Inhibition of TYK2 and JAK1 for the Treatment of Autoimmune Diseases: Discovery of ((S)-2,2-Difluorocyclopropyl)((1R,5S)-3-(2-((1-methyl-1H-pyrazol-4-yl)amino)pyrimidin-4-yl)-3,8-diazabicyclo[3.2.1]octan-8-yl)methanone (PF-06700841)’, Journal of Medicinal Chemistry, 61(19), pp. 8597–8612. Available at: 10.1021/acs.jmedchem.8b00917.

Fourcade, J. et al. (2018) ‘CD226 opposes TIGIT to disrupt Tregs in melanoma’, JCI Insight, 3(14). Available at: 10.1172/jci.insight.121157.

Fuhrman, C.A. et al. (2015) ‘Divergent phenotypes of human regulatory T cells expressing the receptors TIGIT and CD226’, Journal of immunology (Baltimore, Md. : 1950), 195(1), pp. 145–155. Available at: 10.4049/jimmunol.1402381.

Gerstenberger, B.S. et al. (2020) ‘Discovery of Tyrosine Kinase 2 (TYK2) Inhibitor (PF-06826647) for the Treatment of Autoimmune Diseases’, Journal of Medicinal Chemistry, 63(22), pp. 13561– 13577. Available at: 10.1021/acs.jmedchem.0c00948.

Goldstein, J.D. et al. (2016) ‘Inhibition of the JAK/STAT Signaling Pathway in Regulatory T Cells Reveals a Very Dynamic Regulation of Foxp3 Expression’, PLOS ONE, 11(4), p. e0153682. Available at: 10.1371/journal.pone.0153682.

Gorman, J.A. et al. (2019) ‘The TYK2-P1104A Autoimmune Protective Variant Limits Coordinate Signals Required to Generate Specialized T Cell Subsets’, Frontiers in Immunology, 10. Available at: 10.3389/fimmu.2019.00044.

Guan, Q. (2019) ‘A Comprehensive Review and Update on the Pathogenesis of Inflammatory Bowel Disease’, Journal of Immunology Research, 2019(1), p. 7247238. Available at: 10.1155/2019/7247238.

Halim, L. et al. (2017) ‘An Atlas of Human Regulatory T Helper-like Cells Reveals Features of Th2-like Tregs that Support a Tumorigenic Environment’, Cell Reports, 20(3), pp. 757–770. Available at: 10.1016/j.celrep.2017.06.079.

Hovhannisyan, Z. et al. (2011) ‘Characterization of IL-17-producing regulatory T cells in inflamed intestinal mucosa from patients with inflammatory bowel diseases’, Gastroenterology, 140(3), pp. 957–965. Available at: 10.1053/j.gastro.2010.12.002.

Hu, X. et al. (2021) ‘The JAK/STAT signaling pathway: from bench to clinic’, Signal Transduction and Targeted Therapy, 6(1), pp. 1–33. Available at: 10.1038/s41392-021-00791-1.

Ishizaki, M. et al. (2011) ‘Involvement of Tyrosine Kinase-2 in Both the IL-12/Th1 and IL-23/Th17 Axes In Vivo’, The Journal of Immunology, 187(1), pp. 181–189. Available at: 10.4049/jimmunol.1003244.

Izcue, A. et al. (2008) ‘Interleukin-23 Restrains Regulatory T Cell Activity to Drive T Cell-Dependent Colitis’, Immunity, 28(4), pp. 559–570. Available at: 10.1016/j.immuni.2008.02.019.

Jensen, L.T. et al. (2023) ‘Allosteric TYK2 inhibition: redefining autoimmune disease therapy beyond JAK1-3 inhibitors’, eBioMedicine, 97. Available at: 10.1016/j.ebiom.2023.104840.

Keir, M.E. et al. (2020) ‘The role of IL-22 in intestinal health and disease’, Journal of Experimental Medicine, 217(3), p. e20192195. Available at: 10.1084/jem.20192195.

Keohane, C. et al. (2015) ‘JAK inhibition induces silencing of T Helper cytokine secretion and a profound reduction in T regulatory cells’, British Journal of Haematology, 171(1), pp. 60–73. Available at: 10.1111/bjh.13519.

Komatsu, N. et al. (2014) ‘Pathogenic conversion of Foxp3+ T cells into TH17 cells in autoimmune arthritis’, Nature Medicine, 20(1), pp. 62–68. Available at: 10.1038/nm.3432.

Liu, C. et al. (2021) ‘Discovery of BMS-986202: A Clinical Tyk2 Inhibitor that Binds to Tyk2 JH2’, Journal of Medicinal Chemistry, 64(1), pp. 677–694. Available at: 10.1021/acs.jmedchem.0c01698.

Lui, S.-W. et al. (2023) ‘Predicting the clinical efficacy of JAK inhibitor treatment for patients with rheumatoid arthritis based on Fas+ T cell subsets’, APMIS, 131(9), pp. 498–509. Available at: 10.1111/apm.13341.

Masopust, D. and Soerens, A.G. (2019) ‘Tissue-Resident T Cells and Other Resident Leukocytes’, Annual Review of Immunology, 37(Volume 37, 2019), pp. 521–546. Available at: 10.1146/annurev-immunol-042617-053214.

Massa, M. et al. (2014) ‘Rapid and long-lasting decrease of T-regulatory cells in patients with myelofibrosis treated with ruxolitinib’, Leukemia, 28(2), pp. 449–451. Available at: 10.1038/leu.2013.296.

McClymont, S.A. et al. (2011) ‘Plasticity of Human Regulatory T Cells in Healthy Subjects and Patients with Type 1 Diabetes’, The Journal of Immunology, 186(7), pp. 3918–3926. Available at: 10.4049/jimmunol.1003099.

McMurchy, A.N. and Levings, M.K. (2012) ‘Suppression assays with human T regulatory cells: a technical guide’, European Journal of Immunology, 42(1), pp. 27–34. Available at: 10.1002/eji.201141651.

Mease, P.J. et al. (2022) ‘Efficacy and safety of selective TYK2 inhibitor, deucravacitinib, in a phase II trial in psoriatic arthritis’, Annals of the Rheumatic Diseases, 81(6), pp. 815–822. Available at: 10.1136/annrheumdis-2021-221664.

Meyer, A. et al. (2021) ‘Regulatory T cell frequencies in patients with rheumatoid arthritis are increased by conventional and biological DMARDs but not by JAK inhibitors’, Annals of the Rheumatic Diseases, 80(12), pp. e196–e196. Available at: 10.1136/annrheumdis-2019-216576.

Mine, K. et al. (2024) ‘TYK2 signaling promotes the development of autoreactive CD8+ cytotoxic T lymphocytes and type 1 diabetes’, Nature Communications, 15, p. 1337. Available at: 10.1038/s41467-024-45573-9.

Morand, E. et al. (2023) ‘Deucravacitinib, a Tyrosine Kinase 2 Inhibitor, in Systemic Lupus Erythematosus: A Phase II, Randomized, Double-Blind, Placebo-Controlled Trial’, *Arthritis & Rheumatology (Hoboken*, N.J.), 75(2), pp. 242–252. Available at: 10.1002/art.42391.

Morand, E. et al. (2024) ‘TYK2: an emerging therapeutic target in rheumatic disease’, Nature Reviews Rheumatology, 20(4), pp. 232–240. Available at: 10.1038/s41584-024-01093-w.

Moreau, J.M. et al. (2022) ‘Transforming growth factor–β1 in regulatory T cell biology’, Science Immunology, 7(69), p. eabi4613. Available at: 10.1126/sciimmunol.abi4613.

Parampalli Yajnanarayana, S., et al. (2015) ‘JAK1/2 inhibition impairs T cell function in vitro and in patients with myeloproliferative neoplasms’, British Journal of Haematology, 169(6), pp. 824– 833. Available at: 10.1111/bjh.13373.

Pellenz, F.M. et al. (2021) ‘Association of TYK2 polymorphisms with autoimmune diseases: A comprehensive and updated systematic review with meta-analysis’, Genetics and Molecular Biology, 44(2), p. e20200425. Available at: 10.1590/1678-4685-GMB-2020-0425.

Sato, K. et al. (2021) ‘DNAM-1 regulates Foxp3 expression in regulatory T cells by interfering with TIGIT under inflammatory conditions’, Proceedings of the National Academy of Sciences, 118(21), p. e2021309118. Available at: 10.1073/pnas.2021309118.

Satoh-Kanda, Y. et al. (2024) ‘Modifying T cell phenotypes using TYK2 inhibitor and its implications for the treatment of systemic lupus erythematosus’, RMD Open, 10(2), p. e003991. Available at: 10.1136/rmdopen-2023-003991.

Schnell, A., Littman, D.R. and Kuchroo, V.K. (2023) ‘TH17 cell heterogeneity and its role in tissue inflammation’, Nature Immunology, 24(1), pp. 19–29. Available at: 10.1038/s41590-022-01387-9.

Schroeder, M.A. et al. (2020) ‘A phase 1 trial of itacitinib, a selective JAK1 inhibitor, in patients with acute graft-versus-host disease’, Blood Advances, 4(8), pp. 1656–1669. Available at: 10.1182/bloodadvances.2019001043.

Shawky, A.M. et al. (2022) ‘A Comprehensive Overview of Globally Approved JAK Inhibitors’, Pharmaceutics, 14(5), p. 1001. Available at: 10.3390/pharmaceutics14051001.

Strober, B. et al. (2023) ‘Deucravacitinib versus placebo and apremilast in moderate to severe plaque psoriasis: Efficacy and safety results from the 52-week, randomized, double-blinded, phase 3 Program fOr Evaluation of TYK2 inhibitor psoriasis second trial’, Journal of the American Academy of Dermatology, 88(1), pp. 40–51. Available at: 10.1016/j.jaad.2022.08.061.

Sun, L. et al. (2023) ‘T cells in health and disease’, Signal Transduction and Targeted Therapy, 8(1), pp. 1–50. Available at: 10.1038/s41392-023-01471-y.

Thirawatananond, P. et al. (2023) ‘Treg-Specific CD226 Deletion Reduces Diabetes Incidence in NOD Mice by Improving Regulatory T-Cell Stability’, Diabetes, 72(11), pp. 1629–1640. Available at: 10.2337/db23-0307.

Tuomela, K., Salim, K. and Levings, M.K. (2023) ‘Eras of designer Tregs: Harnessing synthetic biology for immune suppression’, Immunological Reviews, 320(1), pp. 250–267.

Wrobleski, S.T. et al. (2019) ‘Highly Selective Inhibition of Tyrosine Kinase 2 (TYK2) for the Treatment of Autoimmune Diseases: Discovery of the Allosteric Inhibitor BMS-986165’, Journal of Medicinal Chemistry, 62(20), pp. 8973–8995. Available at: 10.1021/acs.jmedchem.9b00444.

Yuan, Q. et al. (2007) ‘CCR4-dependent regulatory T cell function in inflammatory bowel disease’, Journal of Experimental Medicine, 204(6), pp. 1327–1334. Available at: 10.1084/jem.20062076.

Yuan, S. et al. (2023) ‘Mendelian randomization and clinical trial evidence supports TYK2 inhibition as a therapeutic target for autoimmune diseases’, eBioMedicine, 89. Available at: 10.1016/j.ebiom.2023.104488.

