## Supplementary Figures and Tables for "TYK2 inhibition enhances Treg differentiation and function while preventing Th1 and Th17 differentiation"

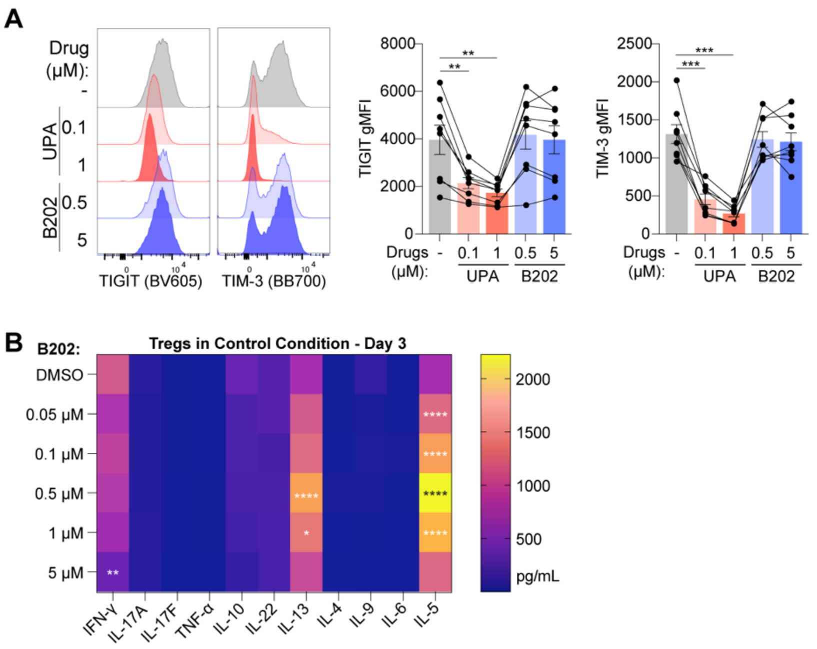


**Supplementary Figure 1. TYK2 inhibition preserves Treg phenotype.** Tregs ($CD4^{+}CD25^{hi}CD127^{lo}$) were isolated from peripheral blood of healthy human donors, stimulated with anti-CD3/CD28, and cultured for 7 days in the presence of BMS-986202 (B202) or upadacitinib (UPA). **(A)** Representative histograms and quantification of TIGIT and TIM-3 gMFI are shown (n=8). **(B)** Supernatant was collected on day 3 of culture and cytokine analysis was carried out (n=5). Statistically significant differences compared to DMSO-treated cells were determined by repeated-measures one-way ANOVA with Dunnett's multiple comparisons test (A) or repeated-measures two-way ANOVA with Dunnett's multiple comparisons test (B). ** p<0.01, *** p<0.001, **** p<0.0001

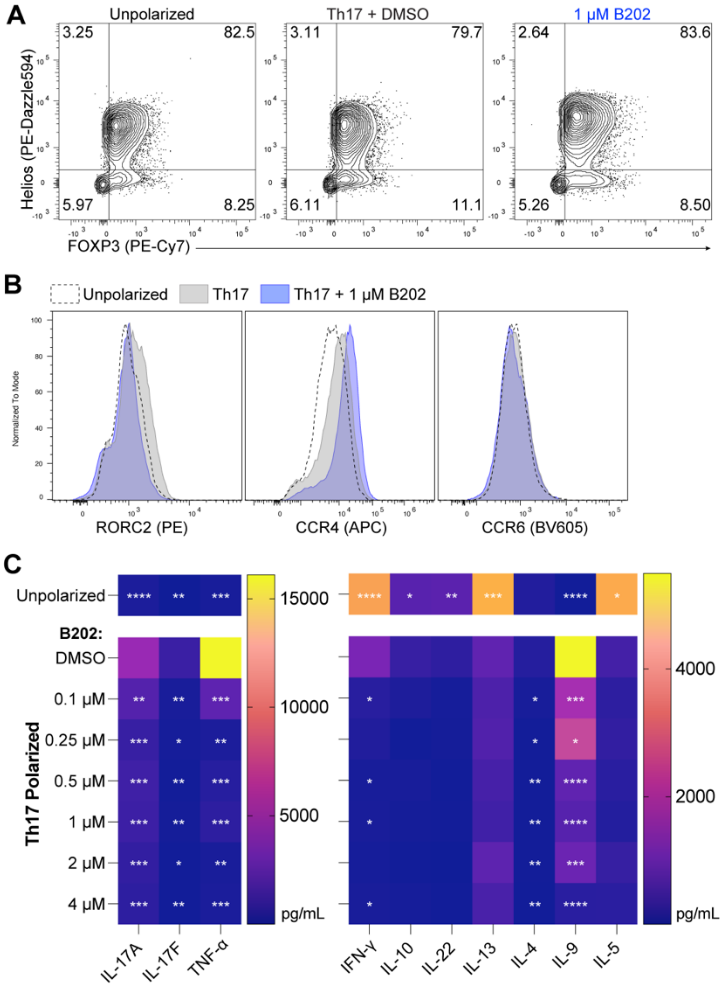


**Supplementary Figure 2. TYK2 inhibition reduces Th17 polarization of Tregs.** Tregs ($CD4^{+}CD25^{hi}CD127^{lo}$) were isolated from peripheral blood of healthy human donors, stimulated with anti-CD3/CD28, and cultured for 7 days in a Th17-polarizing cocktail in the presence of BMS-986202 (B202). **(A)** Representative plots showing expression of FOXP3 and Helios on Tregs after 7 days of culture in different conditions. **(B)** Representative histograms of RORγt, CCR4, and CCR6 expression on Tregs after 7 days of culture in different conditions. **(C)** Tregs were re-stimulated for 24 h and cytokine analysis was carried out on collected supernatant. Statistically significant differences compared to DMSO-treated cells were determined by repeated-measures one-way ANOVA with Dunnett's multiple comparisons test (C). *p<0.05 ** p<0.01, *** p<0.001 **** p<0.0001

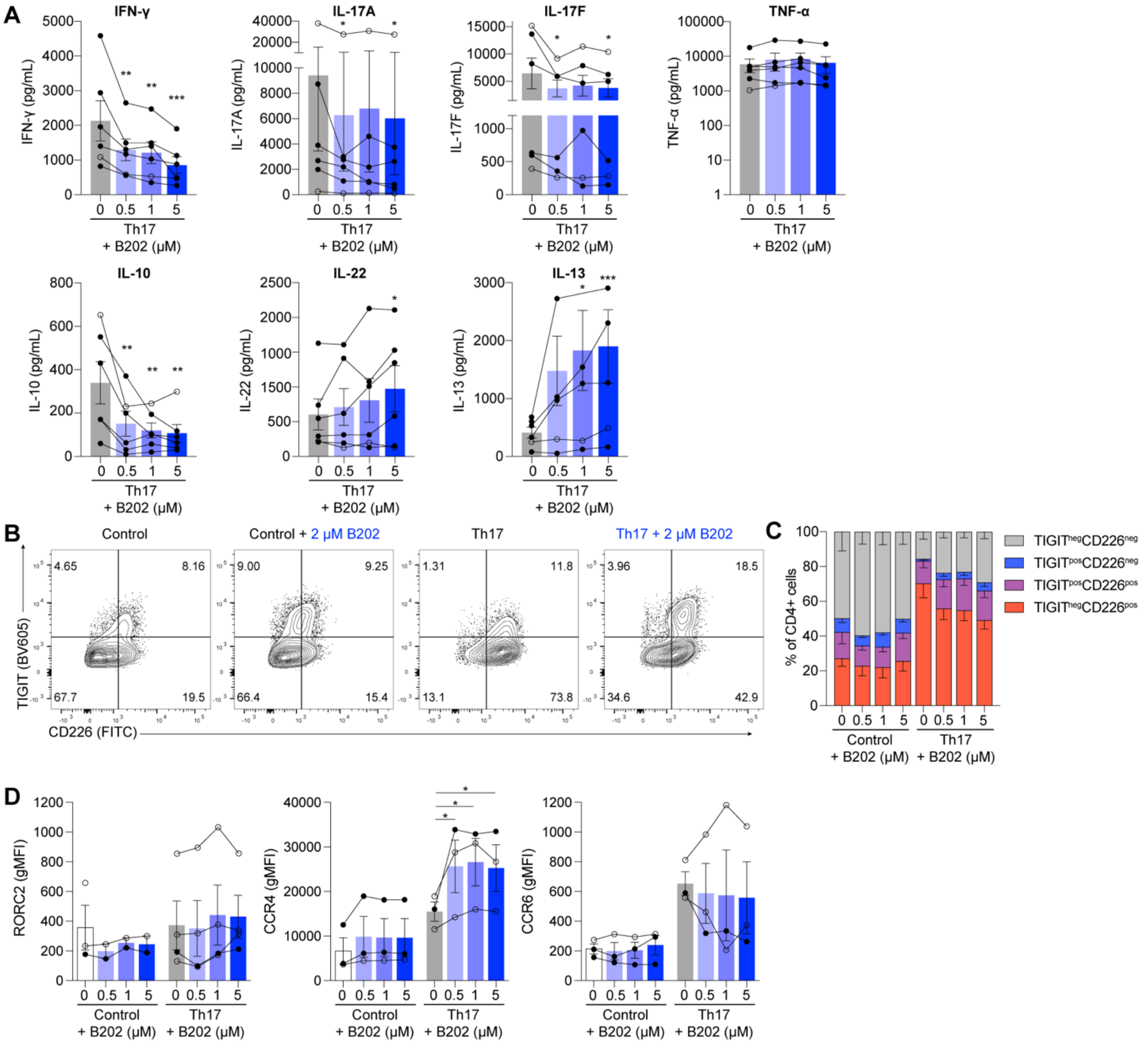


**Supplementary Figure 3. BMS-986202 reduces Th17 polarization of LPMC $CD4^+$ T cells.** Lamina propria mononuclear cells (LPMCs) were isolated from colon biopsies of healthy or IBD patients, stimulated with anti-CD3/CD28 tetramer in the presence of a Th17 cytokine cocktail and BMS-986202 (B202), and cultured for 7 days. **(A)** Cytokine content with supernatant collected after 3 days of culture. Matched data with Figure 6B. **(B)** Representative histogram of CD226 and TIGIT expression on LPMC $CD4^+$ T cells. **(C)** Proportion of $TIGIT^-CD226^-$, $TIGIT^+CD226^-$, $TIGIT^+CD226^+$, and $TIGIT^-CD226^+$ cells of total LPMC $CD4^+$ T cells after 7 days of culture. **(D)** Expression of RORγt, CCR4, and CCR6 on LPMC $CD4^+$ T cells after 7 days of culture. Statistically significant differences compared to DMSO (0 μM)-treated cells were determined by one-way ANOVA with Dunnett's multiple comparisons test. *p<0.05 ** p<0.01, *** p<0.001

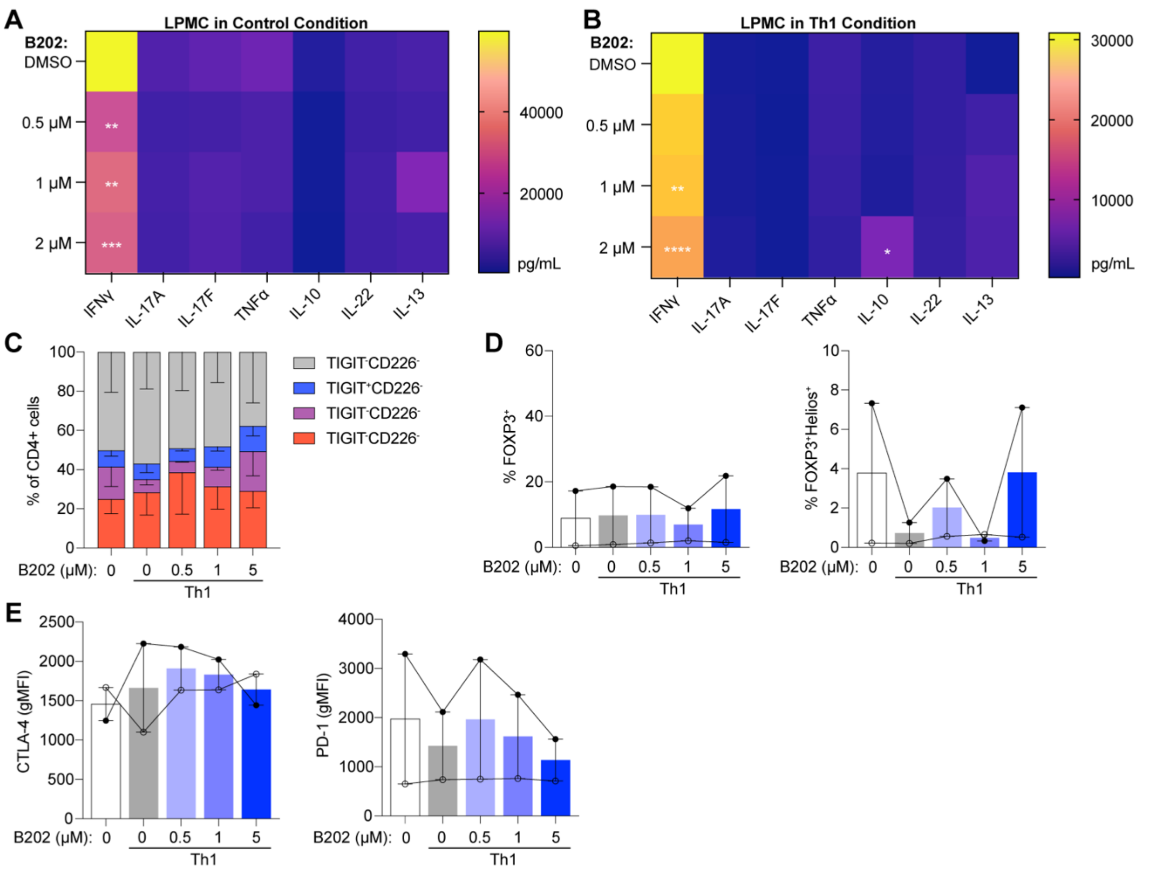


**Supplementary Figure 4. BMS-986202 reduces Th1 polarization of LPMC $CD4^+$ T cells.** Lamina propria mononuclear cells (LPMCs) were isolated from colon biopsies of healthy or IBD patients, stimulated with anti-CD3/CD28 tetramer in the presence of BMS-986202 (B202) and either no exogenous cytokine or with exogenous IL-12, and cultured for 7 days. **(A-B)** Heatmap of cytokine content within supernatant collected after 3 days of culture in no exogenous cytokine (control condition) with varying concentrations of B202. **(B)** Heatmap of cytokine content within supernatant collected after 3 days of culture in exogenous IL-12 to induce Th1 polarization and varying concentrations of B202. **(C)** Proportion of $TIGIT^-CD226^-$, $TIGIT^+CD226^-$, $TIGIT^+CD226^+$, and $TIGIT^-CD226^+$ cells of total LPMC $CD4^+$ T cells after 7 days of culture in Th1-polarizing conditions with or without B202. **(D)** Proportion of $FOXP3^+$ and $FOXP3^+Helios^+$ cells of total LPMC $CD4^+$ T cells after 7 days of culture in Th1-polarizing conditions with or without B202. **(E)** Expression of CTLA-4 and PD-1 on LPMC $CD4^+$ T cells after 7 days of culture in Th1-polarizing conditions with or without B202. Open circles (o) represent healthy patients. Closed circles (•) represent IBD patients. Statistically significant differences compared to DMSO (0 µM)-treated cells were determined by one-way ANOVA with Dunnett's multiple comparisons test. *p<0.05 ** p<0.01, *** p<0.001

**SUPPLEMENTARY TABLES**

### Supplementary Table 1. List of antibodies used in study.

| Antigen | Fluorochrome | Clone | Manufacturer |
|---|---|---|---|
| CCR4 | APC | D8SEE | Invitrogen |
| CCR6 | V650 | G04RE3 | BioLegend |
| CCR6 | BV605 | G04RE3 | BioLegend |
| CD127 | APC-Af700 | R34.34 | Beckman Coulter |
| CD19 | PE-Cy7 | SJ25C1 | Invitrogen |
| CD226 | FITC | TX25 | BD |
| CD25 | BUV395 | 2A3 | BD |
| CD25 | PE | 2A3 | BD |
| CD3 | BUV395 | UCHT1 | BD |
| CD3 | BV605 | UCHT1 | Invitrogen |
| CD39 | BV711 | TU66 | BD |
| CD4 | FITC | RPA-T4 | Invitrogen |
| CD4 | BV480 | SK3 | BD |
| CD4 | BUV496 | SK3 | BD |
| CD4 | PE | OKT4 | Invitrogen |
| CD4 | V500 | RPA-T4 | BD |
| CD45 | BUV805 | HI30 | BD |
| CD45RA | PE-Cy7 | HI100 | Invitrogen |
| CD45RO | eF450 | UCHL1 | Invitrogen |
| CD62L | PerCP-eF710 | DREG-56 | Invitrogen |
| CD8 | BUV563 | RPA-T8 | BD |
| CD80 | BB515 | L307.4 | BD |
| CD86 | PE | IT2.2 | BD |
| CD8a | BV711 | RPA-T8 | BD |
| CTLA-4 | APC | BNI3 | BD |
| CTLA-4 | BV785 | BNI3 | BD |
| CXCR3 | BV421 | G025H7 | BioLegend |
| CXCR3 | BV605 | G025H7 | BioLegend |
| FOXP3 | PE-Cy7 | 236A/E7 | Invitrogen |
| GITR | BV421 | V27-580 | BD |
| Helios | PE-Dazzle594 | 22F6 | BioLegend |
| IFNg | FITC | 4S.B3 | Invitrogen |
| IFNg | BB700 | B27 | BD |
| IL17A/F | BV786 | N49-653 | BD |
| Ki67 | FITC | 20Raj1 | Invitrogen |
| LAP | PE | FNLAP | Invitrogen |
| PD1 | BUV737 | EH12.1 | BD |
| pSTAT1 (pY701) | PE | 4a | BD |
| pSTAT3 (pY705) | APC | 4/P-STAT3 | BD |
| pSTAT4 (pY693) | APC | 38/p-STAT4 | BD |
| pSTAT5 (pY694) | AF488 | 47/pSTAT5 | BD |
| RORC2 | PE | Q21-559 | BD |
| TBET | BV711 | O4-46 | BD |
| TIGIT | BV605 | 741182 | BD |
| TIGIT | PE-Cy7 | MBSA43 | Invitrogen |
| TIM-3 | BB700 | 344823 | BD |

**Supplementary Table 2 Clinical features of IBD and non-IBD LPMC donors.**

| Patient ID | Age | Sex | Smoking Status | IBD Diagnosis | Current Medications |
|---|---|---|---|---|---|
| BMS-2-40 | 39 | M | No | None | N/A |
| BMS-2-41 | 54 | M | No | None | N/A |
| BMS-2-42 | 45 | F | No | None | N/A |
| BMS-3-42 | 31 | F | No | UC | Tofacitinib |
| BMS-3-43 | 40 | M | Yes | CD | Inflectra, Azathioprine |
| BMS-3-44 | 37 | M | No | UC | Infliximab |
| BMS-3-45 | 44 | M | No | CD | Risankizumab |
| BMS-3-46 | 27 | F | No | UC | Mesalamine |
| BMS-3-47 | 34 | F | No | UC | Adalimumab |
| BMS-3-48 | 61 | M | No | UC | Infliximab, Mezavant |
| BMS-3-49 | 26 | M | No | UC | Prednisone |
| BMS-3-51 | 29 | M | No | CD | None |
